# The Liver is an Inflammatory Mediator of Pulmonary Arterial Hypertension

**DOI:** 10.1101/2025.10.03.680270

**Authors:** Navneet Singh, Jordan Lawson, Ashok Ragavendran, Somanshu Banerjee, Andy Hon, Alejandro Vega, Jason Hong, Christopher J. Mullin, Mandy Pereira, Allyson Sherman-Roe, Alexander T. Jorrin, Tiffaney Cayton, Gregory Fishbein, James R. Klinger, William Oldham, Zhiyu Dai, Michael Fallon, Elizabeth O. Harrington, Olin D. Liang, Soban Umar, Corey E. Ventetuolo

## Abstract

The liver’s contribution to pulmonary arterial hypertension (PAH) pathogenesis remains unclear. We hypothesized that the liver promotes inflammatory injury to the pulmonary endothelium. PAH patients without liver disease with pulmonary artery endothelial cell (PAEC) biopsies were included. Liver serologies and imaging were analyzed by unsupervised classification and regression tree (CART) to identify subclinical liver dysfunction clusters. Two machine-learning models predicted cluster assignment and informed differential expression. PAEC transcriptomes were compared to liver and lung data from monocrotaline and Sugen-Hypoxia rats. Liver fibrosis was assessed in rat and human PAH livers. Among 25 PAH patients (76% female, median age 61 [30 – 84] years), CART identified clusters distinguished by Model for End-Stage Liver Disease Sodium (MELD-Na) ≥12, predicting higher pulmonary vascular resistance (ß=0.5 Wood units per point increase in MELD-Na, 95% CI 0.2-0.8, p=0.005) after adjustment for right atrial pressure. Subjects with MELD-Na ≥12 had decreased 6-minute walk distance (353 [120 – 576] m vs. 411[300 – 600] m, p=0.03), with upregulation of apelin, beta-catenin, and immune signaling. Rat lung ECs demonstrated survival and hepatic growth-factor signaling, while rat livers showed immune activation. Rat (20.8 vs 16.6 % area stained, p=0.09) and human PAH livers revealed fibrosis despite absent right ventricular failure, supporting a pathogenic lung-liver axis in PAH.

## Introduction

Pulmonary arterial hypertension (PAH) is a progressive and fatal disorder that can be associated with systemic conditions including liver disease. Hepatic congestopathy is a known complication of right ventricular (RV) dysfunction, and portal hypertension has been linked to long-term prostanoid treatment in PAH.^1^ However, recent work suggests that distinct liver injury phenotypes occur in well-controlled PAH independent of RV failure and without primary liver disease.^2^ While PAH is increasingly recognized as a multisystem disease, the degree to which pulmonary vascular remodeling is perpetuated by inter-organ crosstalk in PAH is unknown.^3–9^

The liver has several key functions as an immune organ, including the presence of resident macrophages (Kupffer cells), induction of antigen specific tolerance, and local surveillance of gut bacteria.^10^ Immunologic and vasoactive factors that first pass through hepatic metabolism^11^ and vascular growth factors secreted by the liver (e.g., bone morphogenetic protein [BMP] 9, a central signaling ligand in PAH) may influence distal vascular beds. Hepatic endothelial cells (ECs) secrete cytokines that contribute to left heart failure via coronary EC dysfunction,^12^ suggesting the liver may function as an endocrine organ via EC crosstalk. Despite the liver’s role as a regulator of immunity and clinically apparent lung-liver phenotypes in PAH, it’s contribution as a driver of pulmonary vascular disease remains unclear.

The current study sought to characterize hepatic involvement in experimental and human PAH without known liver disease. We hypothesized that the liver potentiates systemic inflammation with distal effects on the pulmonary endothelium in PAH. We tested this hypothesis by first using an unsupervised approach to identify subclinical liver dysfunction in PAH patients as determined by laboratory values and Model for End-Stage Liver Disease with Sodium (MELD) scores, which has been applied in non-cirrhotic states to predict outcomes in advanced heart failure, for example.^13^ We then determined the association of cluster membership with pulmonary vascular resistance (PVR) and the differences in differential gene expression in human PAH pulmonary artery endothelial cells (PAECs)^14–16^ between the clusters. We compared these results to lung EC gene expression from two experimental pulmonary hypertension (PH) rat models. Finally, we evaluated bulk RNA sequencing of PH rat livers and assessed the degree of hepatic fibrosis both in rats and patients with PAH.

## Methods

### Sex as a Biological Variable

Both male and female human participants were included, and sexually dimorphic results are reported. Our study examined male rats only. It is unknown whether the findings are relevant for female rats.

### PAH Cohort

Participants with World Symposium on PH Group 1 PAH as diagnosed by a clinical PH provider and PAEC transcriptomic data were included. The clinical diagnosis of PAH was confirmed independently by author N.S. We excluded participants with portopulmonary hypertension (PoPH). All available clinical data (laboratory, imaging, diagnostic codes) were reviewed to confirm the absence of known liver disease defined by elevated transaminases, evidence of synthetic liver dysfunction (elevated international normalized ratio [INR], hypoalbuminemia, thrombocytopenia), or imaging abnormalities including cirrhotic morphology, hepatosplenomegaly or steatosis. The final study cohort consisted of 25 PAH participants (Supplemental Figure 1).

### Human PAEC Biopsies and RNA Sequencing

PAECs were collected at the time of clinically indicated right heart catheterization (RHC) using our previously published cell biopsy method ^14–16^. PAECs were cultured from RHC balloon tips, expanded to passage 3 or 4 and submitted for library preparation and bulk RNA sequencing (Azenta, Cambridge, MA). Libraries were sequenced using a 2 x 50 bp paired-end rapid run on the Illumina HiSeq2500 platform. FastQ files were aligned to the human genome reference sequence GRCh38 using HISAT2 ^17^ and gene IDs were resolved using annotateMyIDs ^18^.

### Cluster Analysis of Clinical Data and Differential Gene Expression Among Clusters

The medical record was retrospectively reviewed, and data including invasive hemodynamics, all available liver imaging and laboratory data (liver transaminases, albumin, platelets, total bilirubin, sodium, creatinine and INR) were collected. The MELD sodium (MELD-Na) 3.0 ^19^ and MELD Xi ^20^ were calculated and included as independent clinical variables. Clinical data and laboratory values were collected as close as possible to the date of PAEC biopsy and within six months of biopsy/RHC catheterization date. An unsupervised classification and regression tree (CART) ^21^ was used to identify distinguishable clusters within the data. Linear regression was subsequently used to model the relationship between these CART-derived clusters and PVR at the time of PAEC cell biopsy, adjusting for right atrial pressure. Linear regression was performed to assess if any of the individual components of MELD-Na contributed to the relationship between PVR and the composite score. Sensitivity analyses were performed to evaluate the influence of connective tissue disease patients and those taking warfarin on the results.

Differentially expressed genes between clusters were generated using linear discriminant analysis (LDA) with principal component analysis (PCA), LDA with weighted gene co-expression analysis (WGCNA), and a random forest model ^22–24^. All three models were assessed for accuracy to predict group assignment. Gene lists generated by all three models were analyzed using Ingenuity Pathway Analysis (Qiagen), significance for pathway enrichment was calculated using a right-tailed Fisher’s Exact Test and gene lists and directional effects predicted using the Ingenuity Knowledge Base.

### Bulk RNA Sequencing of Sugen-Hypoxia and Monocrotaline Livers

To validate findings about hepatic dysfunction in human PAH, we next turned to two leading animal models of PH, the Sugen-hypoxia (SuHx) and monocrotaline (MCT) rat models were performed as described extensively in prior publications by our groups ^25,26^. The right lobe of the liver was removed and fixed in formalin, paraffin embedded, sectioned at 5 micrometers and stained with Masson’s trichrome per standard protocol. All images were acquired using a confocal microscope (Nikon) with a minimum of three images acquired from each slide. Fibrosis was quantified using the Otsu algorithm in Fiji (Image J; NIH). Total RNA was isolated from flash frozen livers from SuHx, MCT, and control rats (four each, all male) using Trizol (Invitrogen). RNA samples were submitted for bulk RNA sequencing to the sequencing core of the University of California, Los Angeles (UCLA); differentially expressed gene lists were generated using *DESeq* in R^27^ and then analyzed using Ingenuity Pathway Analysis (Qiagen) where the significance of enrichment for pathways was calculated using a right-tailed Fisher’s Exact Test.

### Single-cell RNA Sequencing of Sugen-Hypoxia and Monocrotaline Lungs

To further validate our human PAEC findings, we utilized a publicly available data set of single-cell RNA sequencing of SuHX, MCT and control rat lungs ^28^. Raw expression data were normalized, filtered, and clustered using the *Seurat* R package ^29^. Cell types were identified using the labels assigned by the study authors. Differential gene expression in endothelial and immune cell populations was determined using the Wilcoxon rank-sum test. Gene expression was subsequently analyzed using Ingenuity Pathway Analysis (Qiagen).

### Human Liver Biopsies and Masson’s Trichrome Staining

To explore if findings from rat livers were consistent with human disease, we completed complimentary histologic analysis of human tissue from PAH and control patients. Human liver tissue was obtained via warm autopsy at UCLA. Formalin-fixed paraffin-embedded 5 micrometer thick human liver sections (connective tissue disease-associated PAH [CTD-APAH] n=3, idiopathic PAH n=1, PoPH n=5, control n=3) were used for Masson’s Trichrome staining following standard protocols and quantified using the same methodology as rat livers. All images were acquired using a confocal microscope (Nikon). A total of five to ten images were acquired from each slide.

### Statistics

Statistical analysis was performed in R (R Core Team 2024), RStudio (RStudio Team 2025), GraphPad Prism 10.6.1 (GraphPad Software) and Ingenuity Pathway Analysis (Qiagen). Linear regression was performed to evaluate the relationship between MELD and its individual components and PVR as described above. Student’s t-test and Kruskal-Wallis test with post-hoc pairwise comparisons were used to compare the percent area stained of Masson’s trichrome in rat and human livers, respectively. Significance for bulk gene expression pathway enrichment was calculated using a right-tailed Fisher’s Exact Test. For single cell data, differential gene expression was determined using the Wilcoxon rank-sum test.

### Study Approval

Collection of PAECs and clinical data was approved by the Institutional Review Board at Brown University Health (IRB# 001218 and 2221835). For human liver tissue, in accordance with 45 CFR 46, acquisition of the tissue did not constitute human subjects research and used deidentified human tissues provided by UCLA Pathology departmental honest broker, the UCLA Translational Pathology Core Lab (IRB# 11-002504). Animal experiments were approved by the Institutional Animal Care and Use Committee at Brown University Health (IACUC# 2207800).

### Data Availability

Raw and processed PAEC bulk RNA sequencing data are available in the Gene Expression Omnibus under accession number GSE243193.

## Results

### PAH Patients with High-MELD Scores Have Worse Clinical Outcomes

Characteristics of the PAH participants with PAECs available are included in **Table 1**. CART analysis identified that MELD-Na was predictive of PVR. A MELD-Na ≥ 12 was associated with a significantly higher PVR after adjustment for right atrial pressure (i.e., degree of hepatic congestion)(ß=0.5 Wood units per one point increase in MELD-Na, 95% CI 0.2-0.8, p=0.005). Nine participants (36%) had a MELD-Na ≥ 12 (High-MELD group) and sixteen (64%) had a MELD-Na < 12 (Low-MELD group). Both groups were similar in age, distribution of race and ethnicity, and renal function. Compared to the High-MELD group, the Low-MELD group had a higher proportion of female patients (94% vs 44%), fewer connective-tissue disease-associated PAH (CTD-APAH) patients (31% vs 44%) and were less likely to be on anticoagulation (0% vs 33%). Linear regression of the individual variables of MELD-Na revealed that no single variable in the composite score (e.g., creatinine, INR) was responsible for the relationship with PVR. There was no difference in body mass index (BMI), and there was no evidence of hepatic steatosis on available liver imaging (n=15) indirectly suggesting that hepatic steatosis was not present in this cohort. The six-minute walk distance (6MWD) was higher in the Low-MELD group compared to the High-MELD group (411 m vs 353 m, p=0.03). Because connective tissue diseases are associated with systemic inflammation^30,31^ and warfarin can increase the INR (a component of MELD-Na), we then performed sensitivity analyses excluding patients with connective tissue disease (n=9) and taking warfarin (n=2) and found the relationship between MELD-Na and PVR was unchanged.

**Table 1.** Participant Characteristics.

|  | **Low-MELD-Na**<br>**(n=16)** | **High-MELD-Na**<br>**(n=9)** | **All PAH**<br>**Participants**<br>**(n=25)** |
| --- | --- | --- | --- |
| Age, years | 52 (30 – 75) | 68 (43 – 84) | 61 (30 – 84) |
| Female sex, n (%) | 15 (94) | 4 (44) | 19 (76) |
| Race, n (%) |  |  |  |
| Black | 1 (6) | 1 (11) | 2 (8) |
| White | 14 (88) | 7 (78) | 21 (84) |
| Asian | 1 (6) | 0 | 1 (4) |
| Other | 0 | 1 (11) | 1 (4) |
| Hispanic Ethnicity, n (%) | 2 (13) | 1 (11) | 3 (12) |
| Body mass index (kg/m<sup>2</sup>) | 31 (16 – 40) | 35 (23 – 45) | 21 (16 – 45) |
| PAH Etiology, n (%) |  |  |  |
| Idiopathic PAH | 5 (31) | 3 (33) | 8 (32) |
| Hereditary PAH | 3 (19) | 0 | 3 (12) |
| Connective tissue disease-APAH | 5 (31) | 4 (44) | 9 (36) |
| HIV-APAH | 1 (6) | 0 | 1 (4) |
| Congenital heart disease-APAH | 2 (13) | 0 | 2 (8) |
| PVOD | 0 | 2 (22) | 2 (8) |
| Functional Class III or IV, n (%) | 3 (30) | 2 (40) | 5 (20) |
|  | *n=10* | *n=5* | *n=15* |
| 6MWD, meters | 411 (300 – 600) | 353 (120 – 576) | 380 (120 – 600) |
| Hemodynamics |  |  |  |
| RAP, mmHg | 9 (3 – 19) | 10 (4 – 15) | 10 (3 – 19) |
| mPAP, mmHg | 41 (22 – 69) | 44 (26 – 62) | 44 (22 – 69) |
| PAWP, mmHg | 13 (6 – 15) | 11 (6 – 17) | 11 (6 – 17) |
| PVR, Wood units | 4.6 (1.2 – 16.8) | 6.7 (2.2 – 19.6) | 5.3 (1.2 – 19.6) |
| CO, L/min | 5.4 (3.3 – 9.9) | 5.3 (1.9 – 11) | 5.3 (1.9 – 11) |
| MELD-Na | 9 (7 – 11) | 15 (12 – 28) | 10 (7 – 28) |
| Creatinine (mg/dL) | 0.78 (0.52 – 1.26) | 0.96 (0.60 – 1.65) | 0.9 (0.52 – 1.65) |
| Anticoagulation, n (%) | 0 (0) | 3 (33) | 3 (33) |

Data are represented as n (%) or median (range). MELD-Na=model for end-stage liver disease with sodium. PAH=pulmonary arterial hypertension. HIV=human immunodeficiency virus.
PVOD=pulmonary veno-occlusive disease. 6MWD=six-minute walk distance. RAP=right atrial pressure. mPAP=mean pulmonary artery pressure. PAWP=pulmonary artery wedge pressure. PVR=pulmonary vascular resistance. CO=cardiac output.

### PAECs from High-MELD Patients Differentially Upregulate Genes Related to Inflammation

We then used three machine learning models (LDA with PCA, LDA with WGCNA, and a random forest model) to determine gene expression that predicted group assignment (High- vs. Low-MELD); this yielded an accuracy of 0.80, 0.72, and 0.60, respectively. To increase rigor, the union of LDA with PCA and LDA with WGCNA was chosen to build a differentially expressed gene list comparing High- vs. Low-MELD clusters **(Fig 1A).** Pathways analysis demonstrated enrichment for pathways related to apelin signaling, YAP/TAZ, as well as immunity and inflammation **(Fig 1B)**. Network analysis and regulatory networks revealed upregulation of pathways related to apelin, beta-catenin (*CTNNB1*), tumor necrosis factor (*TNF*) and *VCAM-1*, as well as cytokine signaling (e.g., IL-6) in High- vs Low-MELD PAECs **(Fig 2A-B).**

**Figure 1.**
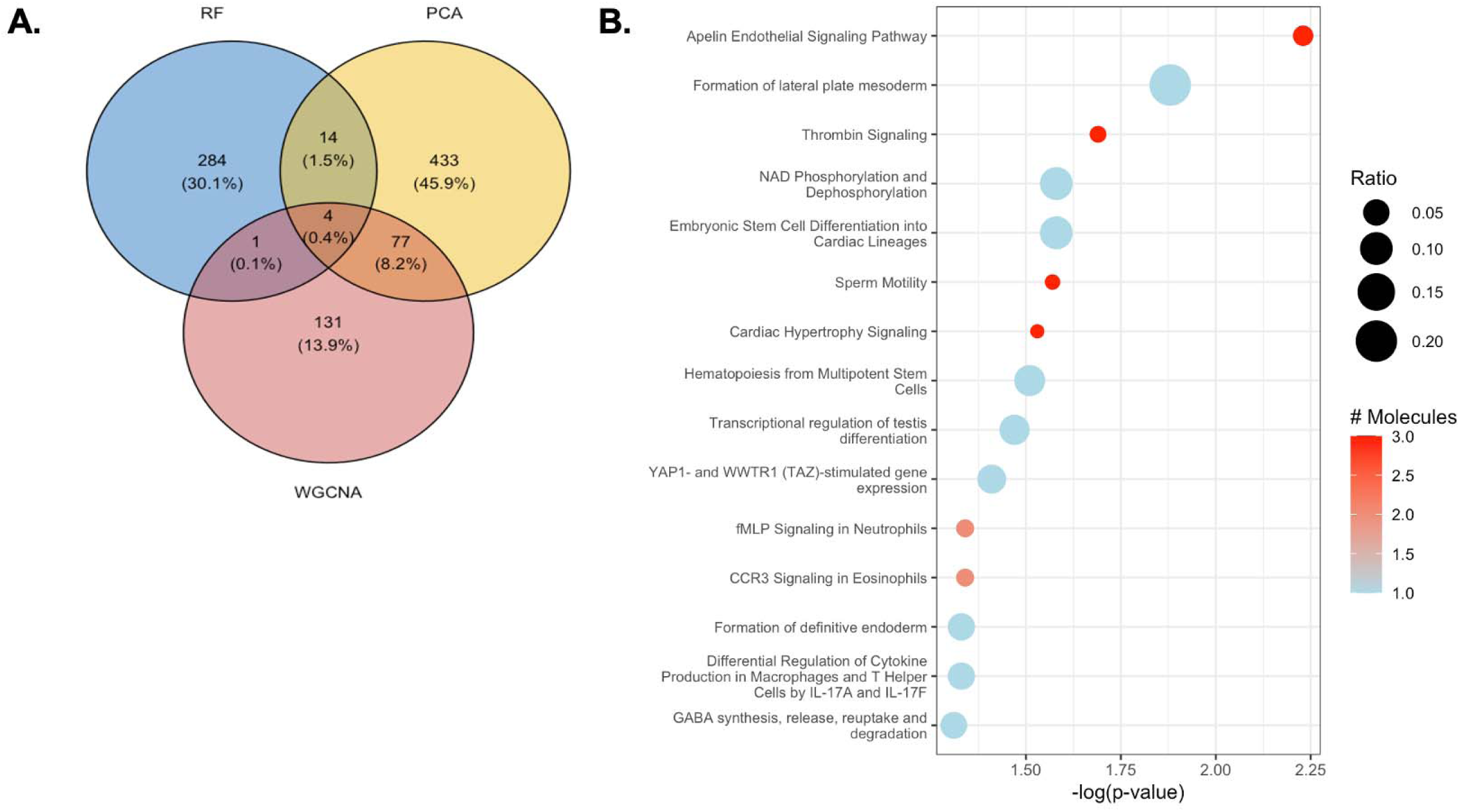
Differences in gene expression between High- and Low-MELD groups were examined using three modeling approaches. **A.** The number and proportion of genes generated by three independent models (linear discriminant analysis (LDA) with principal component analysis (PCA), LDA with weighted gene co-expression analysis (WGCNA), and a random forest (RF) model) and their intersections was reviewed and the intersection of LDA with PCA and WGCNA was used for all subsequent analysis. **B.** Pathways analysis demonstrates enrichment for pathways related to apelin signaling, YAP/TAZ, as well as immunity and inflammation

**Figure 2.**
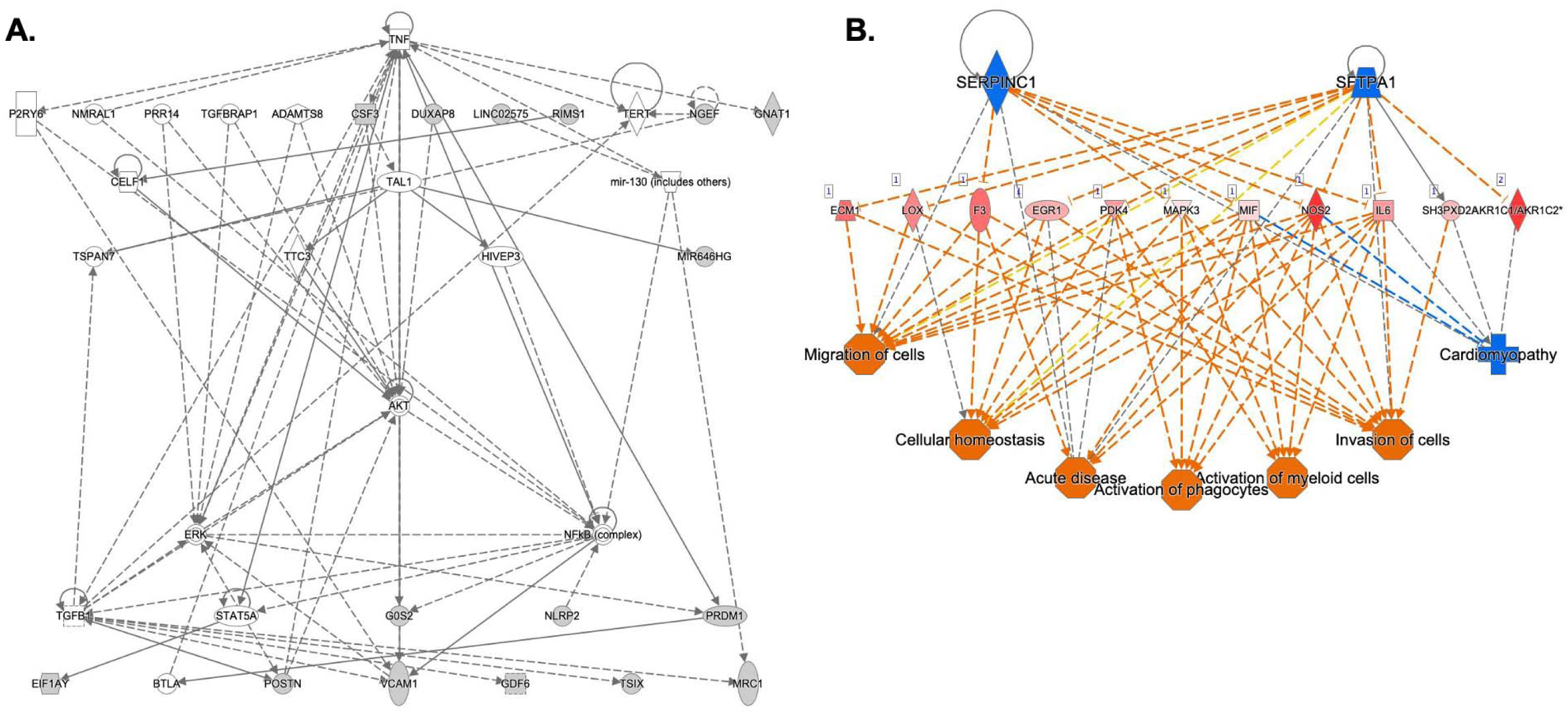
Network analysis of differentially expressed genes in PAECs from High- vs Low-MELD PAH patients reveal High-MELD patients increase expression related to inflammation and immunity. **A.** Network analysis revealed highly relevant expression related to Wnt signaling (*CTNNB1*), inflammation (*TNF*) and recruitment of leukocytes to the pulmonary vasculature (*VCAM1*). **B.** A regulatory network confirmed activation of pathways related to activation of leukocytes and invasion of cells into the pulmonary vasculature mediated by inflammatory signaling (e.g., IL-6), orange=upregulation, blue=downregulation.

### Liver Histology and Gene Expression from Two PH Rat Models Recapitulates Increased Inflammatory Signaling

We then examined whether a similar pro-inflammatory signature existed in the livers of two small animal models of PH. Trichrome staining revealed a trend toward increased perivascular and parenchymal fibrosis in SuHx rat livers compared to controls (20.8 vs 16.6 % area stained, p=0.09)(**Fig 3A-B**). Bulk RNA sequencing of SuHx and MCT rat livers demonstrated similar findings to the High MELD-cluster gene expression from human PAH PAECs. Specifically, we observed increased expression in pathways related to inflammation and TGFβ signaling and decreased expression in pathways related to cellular metabolism. Although the MCT model is known to be characterized by hepatic inflammation and fibrosis,^32^ these themes were recapitulated in the SuHx model (**Fig 4A and B**). A regulatory network demonstrated activation of pathways related to activation and recruitment of leukocytes in these experimental PH livers (**Fig 4C**).

**Figure 3.**
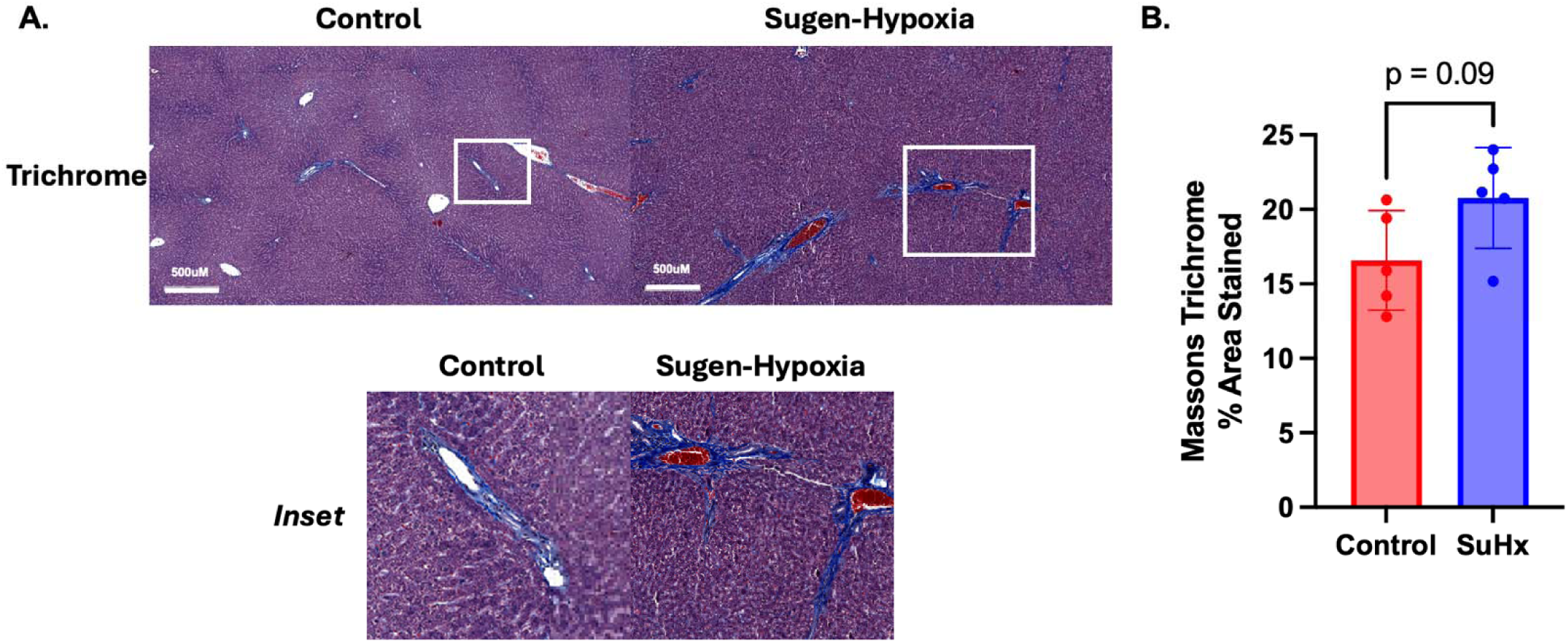
Increased hepatic fibrosis in SuHx rats compared to controls. Control and SuHx rat livers were stained with Masson’s Trichrome which resulted in increased perivascular and parenchymal fibrosis (**A**), 20.8 vs 16.6 % area stained, p=0.09 (**B**). n=5 rats per group. SuHx=Sugen-hypoxia. All images taken at 4X. Scale bar = 500 micrometers.

**Figure 4.**
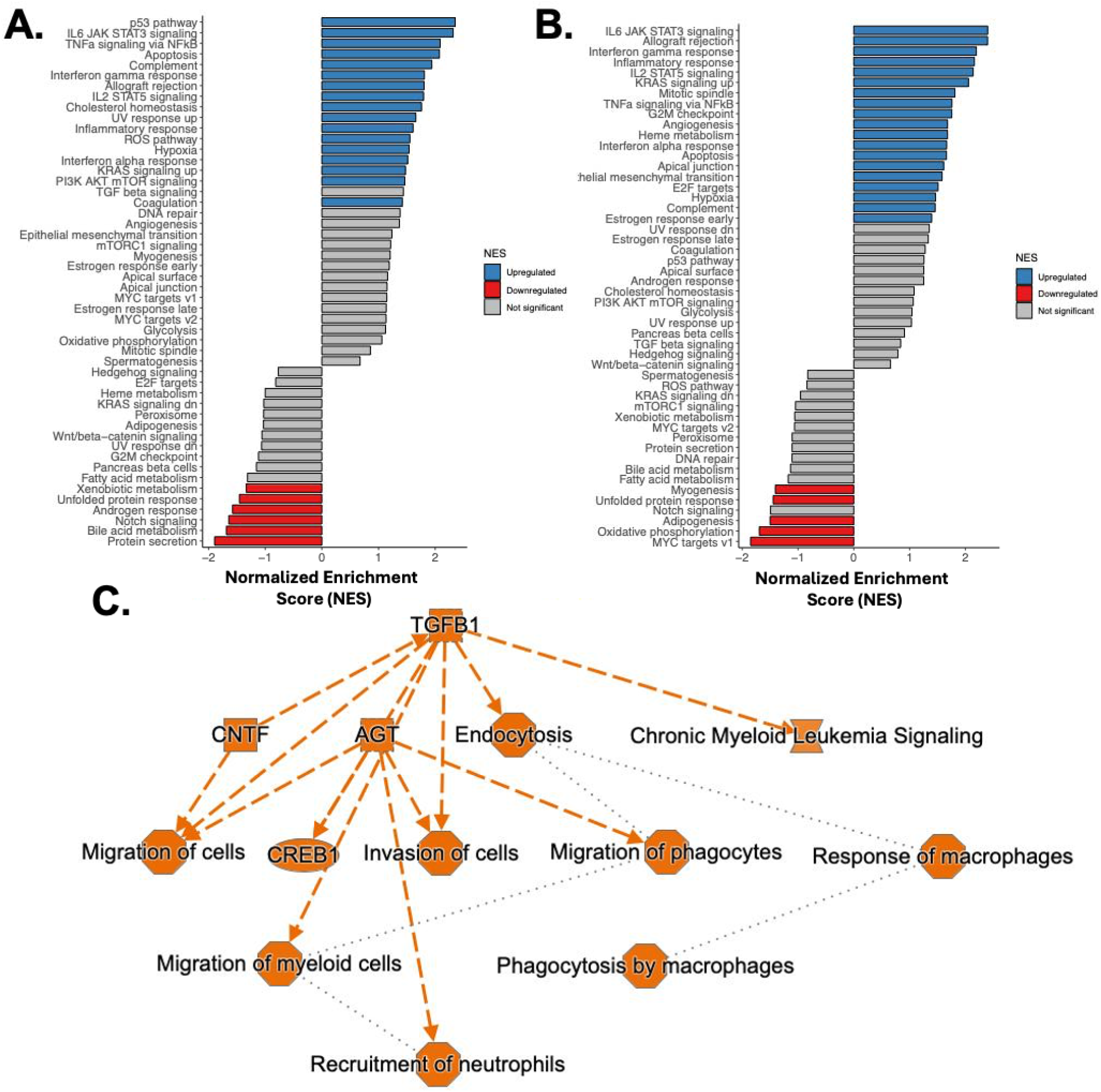
Bulk RNA sequencing of MCT (**A.**) and SuHx (**B.**) male rat livers compared to controls demonstrates increased gene expression in pathways related to inflammation and TGB beta signaling and decreased expression in pathways related to cellular metabolism. **C.** A regulatory network built from differentially expressed genes in both MCT and SuHx rat livers demonstrates activation of pathways related to activation and recruitment of leukocytes.

To understand if our observations in human lungs were consistent with the known biology of these two animal models, differential gene expression in human PAH PAECs from the High-MELD cluster and single cell sequencing data from SuHx and MCT rat lung ECs were then compared. There were 775 genes overlapping in both data sets (**Fig 5A**). Three genes were significantly expressed in both humans and animal regulatory networks: early growth response 1 (*EGR1*), extracellular matrix protein 1 (*ECM1*), and macrophage inhibitory factor (*MIF*). The union of these data sets represented genes with high relevance to cancer biology (**Fig 5B**).

**Figure 5.**
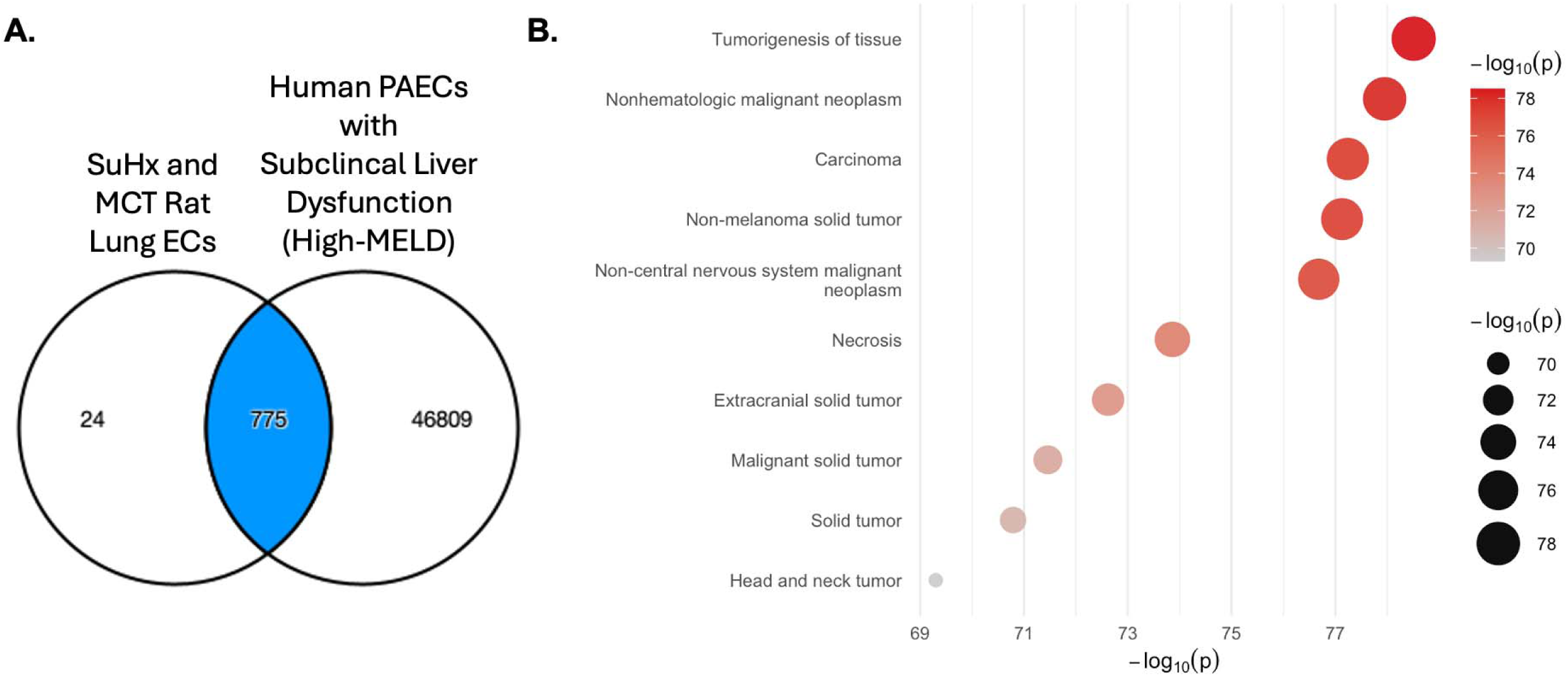
Differential gene expression of SuHx and MCT rat lung ECs demonstrated overlap with PAECs from humans with high-MELD scores (**A**). Genes clustered into highly relevant pathways of cancer biology (**B**). SuHx=Sugen-Hypoxia. MCT=monocrotaline. EC=endothelial cell. PAEC=pulmonary artery endothelial cell.

### Human PAH Livers Demonstrate Increased Fibrosis

Finally, we examined human liver tissue from 12 individuals to determine if the hepatic fibrosis seen in SuHx livers was also present in PAH patients with and without liver disease as compared to controls. Perivascular and parenchymal fibrosis by trichrome staining differed across the three groups (H=7.395, p=0.01). Five PAH patients with liver disease (PoPH) had significantly fibrosis as compared to three control patients (56.3 vs 37.0%, p=0.03). Fibrosis levels in four PAH patients (n=3 CTD-APAH, n=1 idiopathic PAH) without liver disease and without clinical evidence of RV failure (as evidenced by echocardiogram or invasive hemodynamics) had more fibrosis quantitatively and were intermediate between controls and PoPH patients, but these differences were not statistically significant (**Fig 6**).

**Figure 6.**
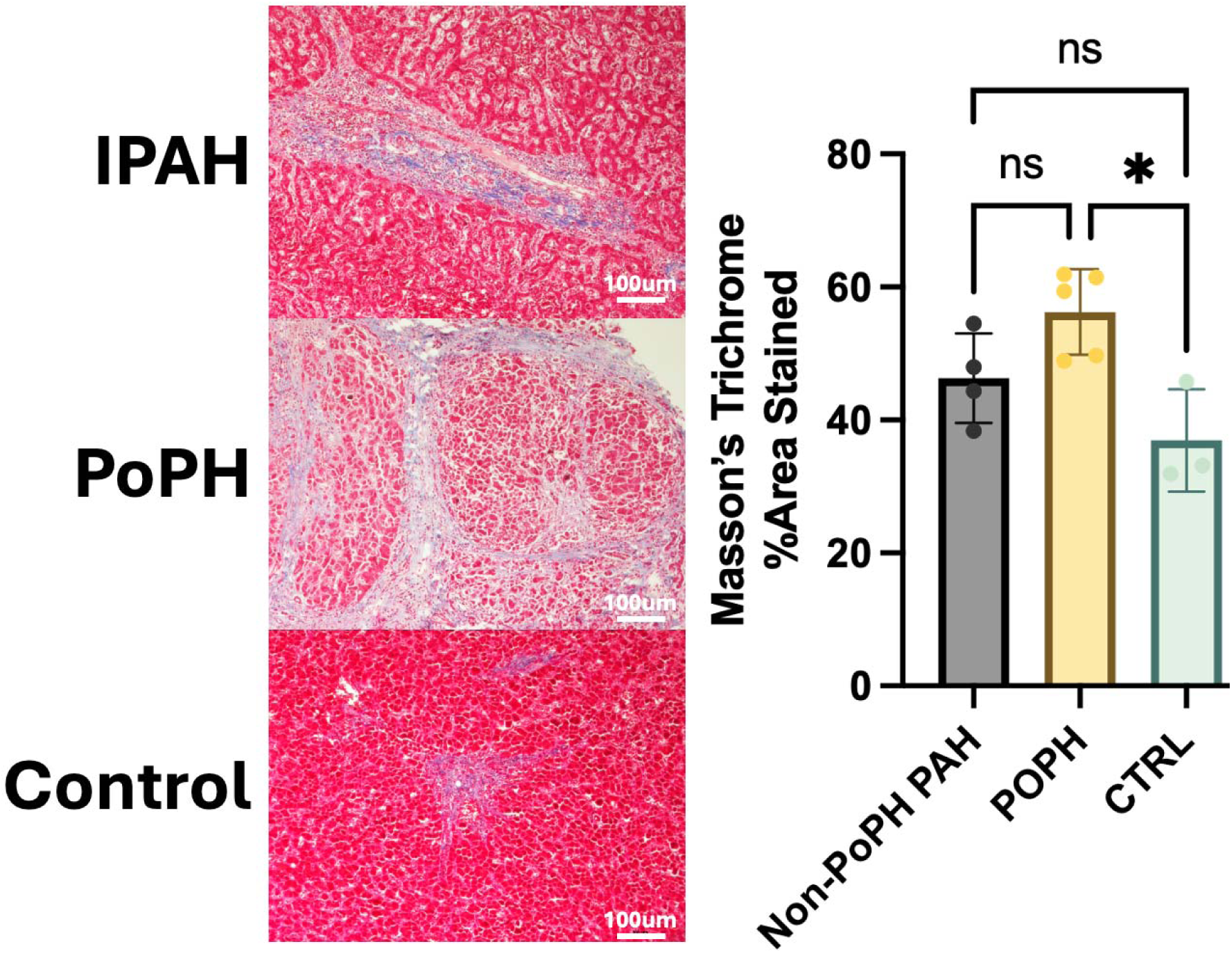
Masson’s trichrome staining of liver tissue from humans with and without pulmonary arterial hypertension (PAH) demonstrate that livers from PAH patients without liver disease suggest an intermediate fibrotic phenotype between those with liver disease (portopulmonary hypertension [PoPH]) and controls (CTRL). IPAH=idiopathic pulmonary arterial hypertension. All images taken at 10x. Scale bar = 100 micrometers. *p=0.03, ns=not significant.

## Discussion

We demonstrate evidence of a lung-liver axis in PAH independent of primary liver disease in both human PAH tissues and two small animal models. Specifically, in an unsupervised cluster analysis, the MELD-Na score was predictive of PVR among PAH patients with no clinical liver disease and independent of elevated right atrial pressure. In PAECs from patients in the High-MELD cluster, i.e., those with subclinical liver dysfunction, there was upregulation in gene expression pathways related to inflammatory activation of the pulmonary endothelium. These findings were recapitulated in two experimental models of PH, one with known liver injury (MCT) but also in SuHx, a model not known to cause direct hepatic injury. There were concordant findings from human PAH PAECs with High-MELD and the two experimental PH models with upregulation of cell survival genes and those related to hepatic growth factor signaling (*EGR1*). Finally, human liver tissue from well characterized PAH patients without liver disease or longstanding RV failure exhibited a pattern of fibrosis that was intermediate between controls and PoPH, mirroring the pattern seen in SuHx rats.

These findings reinforce prior work demonstrating subclinical liver injury predicts clinical outcomes in PAH patients who have participated in clinical trials.^2^ In this report, cholestatic liver injury identified with routine laboratory monitoring predicted worse outcomes, suggesting that bile acids and their metabolism play a role in lung-liver communication in PAH. Circulating bile acids may influence pulmonary endothelial and lysosomal activity and cellular metabolism via the nuclear receptor coactivator 7 (NCOA7).^33^ In the MCT PH model, ketone metabolism is impaired in the liver and leads to activation of the NLRP3 inflammasome suggesting a link between aberrant hepatic metabolism and activation of a systemic inflammatory process,^34^ in-line with our observations in both human and animal tissues. Taken together, these data reinforce our hypothesis that the liver influences the pulmonary circulation as part of a feed-forward loop before (and independent of) underlying liver disease and chronic hepatic congestion.

While specific mechanisms that link the hepatic and pulmonary vascular beds remain elusive, circulating immunologic and vasoactive factors that pass first through (or escape from) hepatic metabolism^11^ and vascular growth factors secreted by the liver are likely important. In children with congenital heart disease, creation of a cavopulmonary shunt causes pulmonary arteriovascular malformations.^35,36^ Circulating BMPs, a family of key signaling ligands in PAH^37^ produced in the liver, are expressed at lower levels in patients with hepatopulmonary syndrome and portopulmonary hypertension as compared to liver disease controls.^38,39^ We noted upregulation of apelin signaling in our PAECs, which we speculate may be a protective compensatory response as apelin is upstream of BMP signaling, including BMPR2.^40^

In human PAH PAECs with High-MELD and PH rat lungs, we noted increased expression of *EGR1*. EGR1 is a transcriptional regulator which plays a role in cell survival, proliferation, and death. Hepatocyte growth factor (HGF) can induce the expression of *EGR1* to regulate these processes^41,42^ and in hepatocellular cancer, EGR1 is implicated in malignant cells’ escape from anticancer drugs via stabilization of microtubules and autophagy.^43,44^ Increased HGF is associated with worse survival in PAH,^45^ however HGF supplementation has been shown to be beneficial in experimental PH.^46^ Hepatokines may modulate endothelial dysfunction in PAH but require additional investigation. We also noted upregulation of gene expression related to cell proliferation (*ECM1*)^47^ and escape from cell death via Wnt signaling (*CTNNB1*),^48,49^ reinforcing well established paradigms of PAEC dysregulation in PAH.^50^

Perivascular CD68^+^ macrophages and other inflammatory cells are prominent in plexiform lesions in both animal models and humans with PAH.^51–54^ Macrophages are differentially polarized in PAH,^55,56^ and depletion or inactivation of macrophages can prevent disease, including in portopulmonary hypertension.^57–61^ PAECs are known to recruit leukocytes via increased expression of VCAM-1, ICAM-1 and E-selectin.^62^ Here, we demonstrated increased expression of *VCAM-1* in the PAECs of High-MELD PAH patients.^63,64^ Finally, cytokine signaling (e.g., IL-6, TNF) was upregulated in High-MELD PAECs concordant with established literature that circulating inflammatory cytokines and chemokines are elevated in idiopathic PAH and some, specifically IL-6 and TNF-α, correlate with outcomes.^65,66^

This study has limitations. We cannot definitively exclude hepatic congestion as contributing to our findings, although we adjusted for right atrial pressure in our analysis to minimize confounding and echocardiogram and invasive hemodynamics were used to exclude RV dysfunction in patients with liver biopsies. We acknowledge that the MELD-Na was not developed for use in a population without liver disease nor does it comprehensively capture liver function, however it has been used in to predict outcomes in left heart failure.^13^ Our results were unchanged when we excluded connective tissue disease participants, who may be more prone to autoimmune liver disease and those taking warfarin, which may pharmacologically increase the INR (a component of MELD-Na). We had similar results when we used alternative MELD scores (MELD-Xi). A sensitivity analysis evaluating individual components of MELD-Na demonstrated that no single factor explained the relationship with PVR. The low MELD-Na cluster was predominantly female (who had higher six-minute walk distances), consistent with the known female advantage in PAH.^67^ Furthermore, we have confirmed the hypotheses generated by our unsupervised analyses in two animal models of PH – one with known liver injury and one without – and human PAH, demonstrating hepatic inflammation and fibrosis is associated with activation of the pulmonary endothelium via expression of *VCAM-1, TNF,* and *MIF.* Whether these observations in SuHx are due to Sugen5416 directly acting on the liver or pulmonary endothelial injury and lung-liver crosstalk is unknown. However, VEGF inhibition in the liver has been shown to be antiproliferative and antifibrotic,^68,69^ whereas we observed increased fibrosis and inflammation in SuHx. Any prior observations of liver injury in SuHx are in chronic RV failure models.^70^ While BMI may be a poor measure of body composition^71^, our conclusion that steatohepatitis is not a confounder in this study is supported by the lack of fatty liver infiltration on all available liver imaging. Human liver tissue samples were limited, but similar themes emerged across human and animal tissues. Failure to detect significant differences between the degree of fibrosis in PAH livers without liver disease (non-PoPH) and control livers may have been due to sample size. While our observations need to be confirmed beyond the transcript level, we contend that consistent observations in humans, two animal models, and across lung and liver tissue sources are compelling.

## Conclusion

In conclusion, we have demonstrated that the liver plays a role in the development of PAH by mediating inflammatory activation of the pulmonary endothelium. This relationship is characterized by hepatic inflammation and fibrosis, even in patients and animal models not known to have detectable liver disease. Taken together, this supports key interorgan modulation and a lung-liver axis early in the PAH disease continuum.

## Author Contributions

NS and CEV conceptualized the study and drafted the manuscript. NS, CJM, MP, ASR, ATJ, TC, JRK, WO and CEV collected and propagated PAECs. JL and AR performed the cluster analysis of human PAEC bulk RNA sequencing. JH, SB and SU performed and analyzed rat liver bulk RNA sequencing. NS performed rat and analyzed rat and human histology. SB, AH, AV, GF, SU performed human liver histology. Data analysis and interpretation was performed by NS, ODL and CEV. The manuscript was reviewed and approved by all authors.

## Acknowledgements

The authors would like to acknowledge all the patients who participated in these studies.

