## Supplemental Figure 1 for "The Liver is an Inflammatory Mediator of Pulmonary Arterial Hypertension"

Excluded (n=1)

Portopulmonary Hypertension

Group 1 PAH patients without known liver disease (n=25)

Assessed for eligibility (n=26)

**Supplemental Figure 1.** Flow diagram of patient screening, eligibility and exclusion for the study cohort. Among Group 1 PAH subjects, the diagnoses were independently confirmed and those with underlying liver disease (n=1) were excluded. The final cohort consisted of 25 Group 1 PAH subjects without any clinically detectable liver disease. *PAH=pulmonary arterial hypertension.*
